# Phosphatidylserine-binding receptor, CD300f, on macrophages mediates host invasion of pathogenic and non-pathogenic rickettsiae

**DOI:** 10.1101/2024.05.10.593542

**Authors:** Oliver H. Voss, Imran Moin, Hodalis Gaytan, Mohammad Sadik, Saif Ullah, M. Sayeedur Rahman

## Abstract

Some arthropod-borne obligate intracellular rickettsiae are among the most virulent human pathogens. *Rickettsia* species modulate immune (e.g., macrophages; MΦ) and non-immune cell (e.g., endothelial cells) responses to create a habitable environment for host colonization. In particular, MΦ play a crucial role in either terminating an infection at an early stage or succumbing to bacterial replication and colonization. However, our understanding on how *Rickettsia* species invade host cells, including MΦ, remain poorly defined. In this study, we describe a mechanism of host invasion by *Rickettsia* species, involving rickettsial phosphatidylserine (PS), as a ligand, and the CD300f receptor on MΦ. Using bone marrow-derived macrophages (BMDMΦ) from wild-type (WT) and CD300f^-/-^ mice, we demonstrated that engulfment of both pathogenic *R. typhi* (the etiologic agent of murine typhus) and *R. rickettsii* (the etiologic agent of Rocky Mountain spotted fever) species as well as the non-pathogenic *R. montanensis* was significantly reduced in CD300f^-/-^ BMDMΦ as compared to that of WT BMDMΦ. Furthermore, our mechanistic analysis suggests bacterial PS as the potential source for the CD300f-mediated rickettsiae engulfment by MΦ. *In vivo* infection studies using WT and CD300f^-/-^ C57BL/6J mice showed that CD300f^-/-^ animals were protected against *R. typhi*-or *R. rickettsii*-induced fatal rickettsiosis, which correlated with levels of bacterial burden detected in the spleens of mice. Adoptive transfer studies further revealed that CD300f-expressing MΦ are important mediators to control rickettsiosis *in vivo*. Collectively, our findings describe a previously unappreciated role for the efferocytic receptor, CD300f, to facilitate engulfment of rickettsiae within the host.

## Introduction

Vector-borne diseases represent a serious concern for global public health. In the United States, many vector-borne pathogens, transmitted by hematophagous arthropods (e.g., ticks and fleas), are on the rise, as exemplified by recent outbreaks of *R. rickettsii*, an etiologic agent of Rocky Mountain Spotted Fever, in Arizona (1) and of *R. typhi*, an etiologic agent of murine typhus, in California (2) and in Texas (3). Human infection with rickettsiae occurs via infected arthropods (e.g., ticks and fleas) either through their bite or deposited infected feces onto the host’s dermis and mucosal surfaces. *Rickettsia* species, at the infection sites, encounter tissue-resident immune cells, like macrophages (MΦ) and dendritic cells (4). MΦ play a crucial role in either terminating an infection at an early stage of host invasion, which commonly is the fate of non-pathogenic *Rickettsia* species, or succumbing to bacterial replication and pathogen colonization as well as dissemination to distant organs of host (4). However, various key cellular processes of rickettsial obligate lifestyle, which includes ***I)*** internalization by phagocytosis into professional phagocytes (e.g., MΦ) or endothelial cells, ***II)*** regulation of membrane dynamics and intracellular trafficking, and ***III)*** evasion of host defenses to establish an intracytosolic replication niche, remain poorly defined and have impaired the development of effective interventions against pathogenic rickettsiae (4, 5). Specifically, our understanding on rickettsiae-induced phagocytosis relies mostly on reports from endothelial cells, leaving the mechanism of rickettsiae engulfment by host defense cells, like MΦ, mostly unknown. In fact, preceding research on endothelial cells has revealed the identity of four host receptors Ku70 (6), α2β1 (7), FGFR1 (8), Epac1 (9), with limited effects on *Rickettsia* invasion (∼ 40%) when silenced individually, supporting the hypothesis of an alternative receptor (on host)-ligand (on rickettsiae) system for rickettsial engulfment. Yet the precise mechanism(s) remain to be elucidated.

Phagocytosis of apoptotic cells by MΦ [aka efferocytosis], is critical for maintaining homeostasis by removing apoptotic cells before becoming proinflammatory and immunogenic causing autoimmune diseases (10, 11). Importantly, the same mechanism can also be hijacked by various pathogens, including viruses and bacteria, to promote host cell invasion (12). In fact, pathogens have been shown to exploit efferocytosis through externalization of “eat-me” signals [e.g., phosphatidylserine (PS)] to gain access to various PS-binding receptors on MΦ, including CD300s (12, 13) to evade host immune surveillance, and to promote their dissemination within the host (12). The CD300 family are type I transmembrane cell surface receptors with a single immunoglobulin (Ig)V-like extracellular domain that can transmit either activating or inhibitory signals (11, 14, 15). Specifically, activating receptors have usually short intracellular tails and gain activation potential through binding of adaptor molecules (e.g., DAP12) harboring immunoreceptor tyrosine-based activation motifs. In contrast, inhibitory receptors have immunoreceptor tyrosine-based inhibitory motifs within their intracellular tails. The orthologous mouse family has a variety of names, including CMRF-like molecules (CLM) (15), but for simplicity, herein, we use the human nomenclature for both species. CD300f is predominantly expressed by myeloid cells and is unique as it displays both activating and inhibitory signaling capabilities (11, 15). Moreover, CD300f is a PS-binding receptors, which regulates efferocytosis in professional phagocytes (e.g., MΦ) (11, 15–19). In line with such an unique role, CD300f has also been shown to play an important role in modulating allergic, inflammatory, autoimmune viral and bacterial responses. However, the contributing role of PS-binding receptor CD300f during rickettsiae engulfment in MΦ (or other immune defense cells) causing ultimately fatal rickettsiosis has not been investigated.

Here, we assessed the functional role of CD300f in rickettsial pathogenesis, by first determining the phagocytic capability of bone marrow-derived macrophages (BMDMΦ) from wild-type (WT) and CD300f^-/-^ mice infected with *R. typhi*, *R. rickettsii*, or *R. montanensis*, and showed that engulfment of pathogenic as well as non-pathogenic *Rickettsia* species was reduced in CD300f^-/-^ BMDMΦ when compared to that in WT BMDMΦ. Further mechanistic analysis indicated that CD300f receptor modulates the phagocytosis of all three *Rickettsia* species, likely via rickettsial PS as ligand. *In vivo* infection studies, using our previously established mouse model of fatal rickettsiosis (20), revealed that CD300f^-/-^ mice (17), but not WT animals, were protected against *R. typhi*-or *R. rickettsii*-induced lethality. Using adoptive transfer studies, we also showed that CD300f-expressing MΦ were critical mediators in controlling rickettsial infection *in vivo*. Collectively, our findings describe a previously unappreciated role of CD300f receptor during rickettsial infection.

## Results

### CD300f modulates engulfment of pathogenic and non-pathogenic rickettsiae

Efferocytosis carried out by professional phagocytes, like MΦ, is critical for maintaining cellular homeostasis by removing large quantities of apoptotic cells before they become proinflammatory and immunogenic causing the development of autoimmune diseases (10–12). Intriguingly, the same mechanism can also be hijacked by viruses and bacteria to promote their host invasion by utilizing PS externalization to gain access to PS-receptors, including Tyro3, Axl, MerTK, TIM1/4, or CD300s (e.g., CD300f) (12, 13). Importantly, MΦ are one of the cell types first encountered by rickettsiae at the site of inoculation and are considered key players to either terminate the infection early at the infection site or allow pathogen colonization and subsequent dissemination within the host (4). However, how rickettsiae gain access into host defense cells, like MΦ, remains to be investigated. In this study, we test the hypothesis that *Rickettsia* species utilize the PS-binding receptor CD300f to invade and colonize MΦ. In this effort, we determined the invasion of *R. typhi*, *R. rickettsii*, and *R. montanensis* into BMDMΦ, from either WT or CD300f-deficient mice, by immunofluorescent assay (IFA) (21) and observed that CD300f facilitated the internalization of all tested rickettsiae (Figs. 1A-C). To further confirm these data, we evaluated the bacterial burden (measured by *gltA* expression) by employing RT-qPCR. For all three *Rickettsia* species found similar results at early time points (Figs. 1D-F), however, *R. montanensis* showed a reduced growth at 24 hrs (Fig. 1F), supporting our previous reporting that *R. montanensis*, but not *R. typhi* and *R. rickettsii*, is cleared by MΦ (20, 22). As the CD300 family is composed of several members (11, 15, 17, 18, 23), we determined the expression of various members of CD300 family by RT-qPCR in uninfected WT and CD300f-deficient BMDMΦ. The data showed that the lack of CD300f expression did not affect the expression of other CD300 members, including *CD300a*, *CD300b*, *CD300c*, and *CD300d* (Fig. S1A). Of note, expression level of *CD300c* was significantly lower as compared to all other tested *CD300*s in both WT and CD300f-deficient BMDMΦ (Fig. S1A). Together our data indicate an important role for CD300f by modulating rickettsiae engulfment in MΦ. To further validate this hypothesis, we evaluated the engulfment of all three *Rickettsia* species using BMDMΦ isolated from WT, CD300f^-/-^, and CD300d^-/-^ (a highly expressed activating receptor in MΦ, Fig. S1A) mice by IFA and RT-qPCR. Our data revealed that, unlike in WT or CD300d^-/-^ BMDMΦ, engulfment and bacterial burden of *R. typhi*, *R. rickettsii*, and *R. montanensis* were significantly reduced in CD300f^-/-^ BMDMΦ (Figs. S1B-G) suggesting that CD300f receptor plays a key role in the engulfment of pathogenic and non-pathogenic rickettsiae by MΦ.

**Fig. 1.**
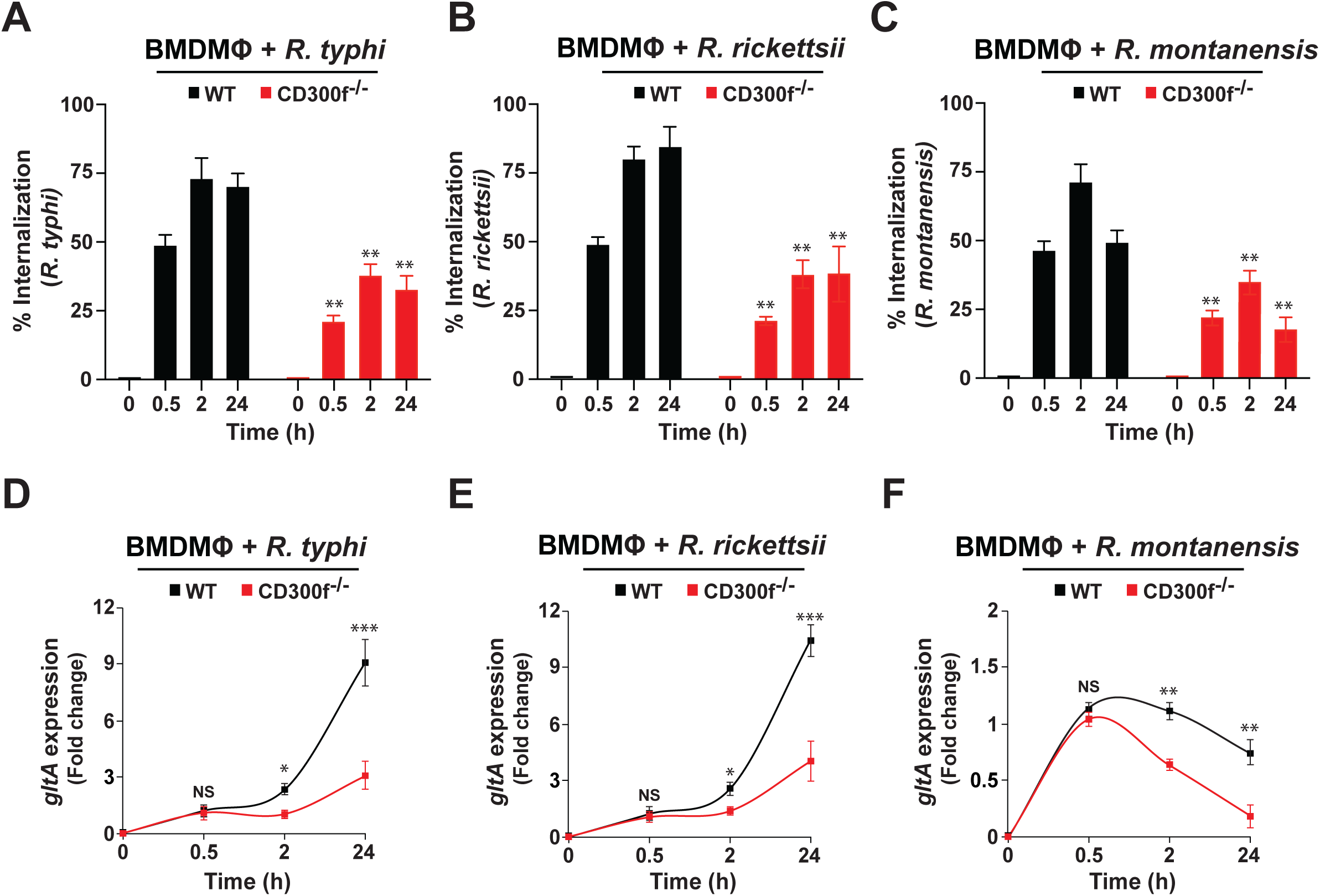
Phosphatidylserine receptor, CD300f, facilitates the engulfment of pathogenic and non-pathogenic rickettsiae. BMDMΦ from WT, or CD300f^-/-^ mice were infected with partially purified *R. typhi*, *R. rickettsii*, and *R. montanensis* at a multiplicity of infection (MOI) of 20 (0.5 and 2 h) and 5 (24 h). (A-C) Rickettsial invasion was monitored by IFA at 0.5, 2, and 24 hours post infection (hpi) as described previously (21). Bacterial burdens of WT and CD300f^-/-^ BMDMΦ infected with *R. typhi* (D), *R. rickettsii* (E), or *R. montanensis* (F) was determined at 0.5, 2, and 24 hpi (n = 5 per group) by RT-qPCR. RCN of *gltA* expression of rickettsiae was normalized by the expression of the housekeeping host gene, *GAPDH*. Error bars in panels A-F represent means ± SEM from five independent experiments. NS, nonsignificant; \**P* ≤ 0.05; \*\**P* ≤ 0.01; \*\*\**P* ≤ 0.005.

### Phosphatidylserine is involved in the CD300f-mediated engulfment of rickettsiae

As CD300f primarily functions as a phosphatidylserine (PS)-binding receptor to regulate phagocytosis events, like efferocytosis (11, 16–19, 23), we sought to test the hypothesis that CD300f-mediated engulfment of *Rickettsia* species by MΦ involves rickettsial PS, as ligand. In this effort, we incubated partially purified *R. typhi*, *R. rickettsii*, and *R. montanensis* with increasing concentrations of recombinant Annexin V (rAnxV), a molecule known to bind to and inhibit PS-mediated processes, like efferocytosis (16, 17, 23). Following incubation, the rAnxV-treated rickettsiae were utilized to infect WT BMDMΦ. Rickettsiae engulfment and burden were evaluated by IFA and RT-qPCR respectively (20, 21). Our data revealed that pretreatment with rAnxV significantly reduced the engulfment (Figs. 2A) and burden (Figs. 2B) of all three rickettsiae in a dose-dependent manner. In addition, experiments using heat-inactivated (HI) rAnxV were employed as specificity control and failed to inhibit the bacterial invasion in MΦ (Figs. 2A-B), indicating that PS is a putative ligand contributing to the rickettsial invasion process.

**Fig. 2.**
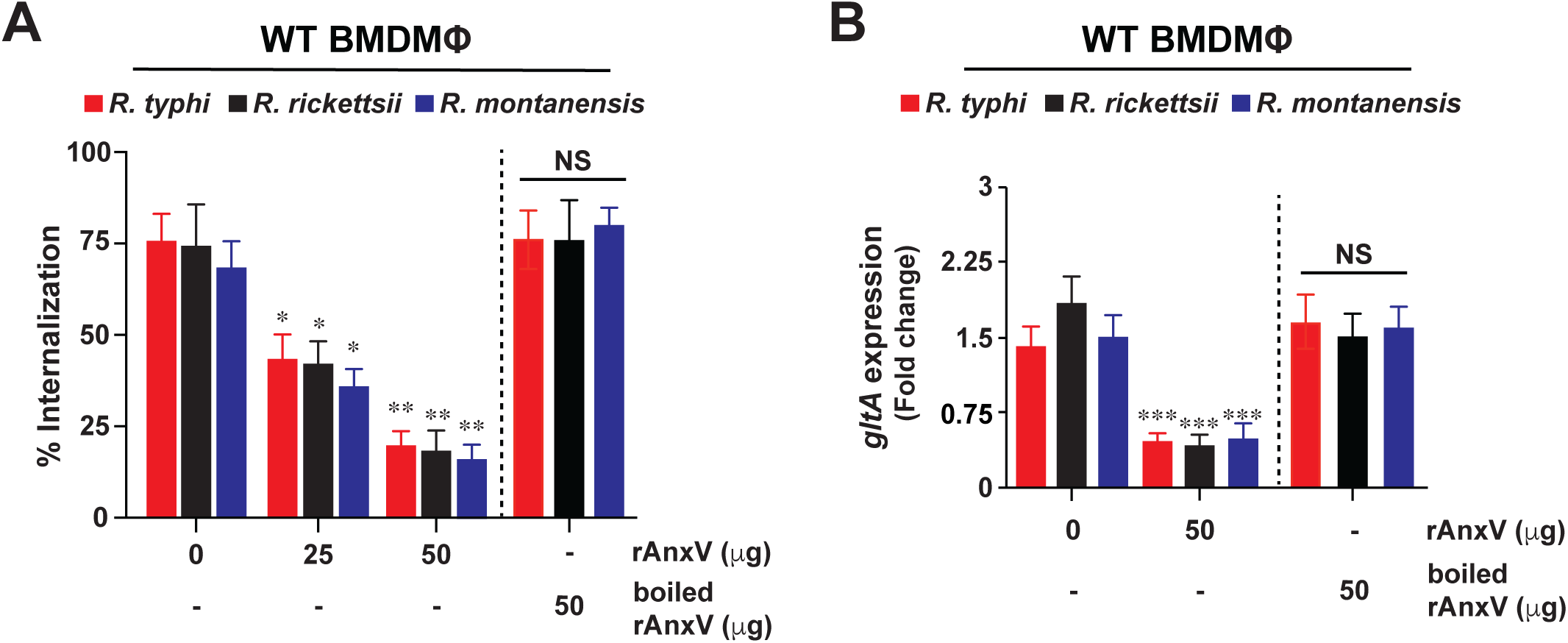
Blocking phosphatidylserine reduces invasion of pathogenic and non-pathogenic rickettsiae in macrophages. Partially purified *R. typhi*, *R. rickettsii*, and *R. montanensis* were pretreated for 0.5 h with various amounts of recombinant Annexin V (rAxnV) and then utilized to infect BMDMΦ from WT mice. Cells were infected with rickettsiae (untreated or pretreated with rAxnV) for 2 hpi using a MOI of 20. Boiled rAnxV served as control. (A) Rickettsial invasion was monitored by IFA at 2 hpi as described previously (21). (B) Bacterial burden of WT BMDMΦ infected with *R. typhi*, *R. rickettsii*, or *R. montanensis* was determined at 2 hpi (n = 5 per group) by RT-qPCR. RCN of *gltA* expression of rickettsiae was normalized by the expression of the housekeeping host gene, *GAPDH*. Error bars in panels A and B represent means ± SEM from five independent experiments. NS, nonsignificant; \**P* ≤ 0.05; \*\**P* ≤ 0.01; \*\*\**P* ≤ 0.005.

### Pathogenic and non-pathogenic *Rickettsia* species express phosphatidylserine

To further elucidate the mechanism of PS-CD300f-mediated engulfment of rickettsiae, we wanted to delineate the source of PS. Detection of PS using fluorescently labeled AnxV or anti-PS antibodies are commonly used to evaluate apoptotic events in eukaryotic cells, however only few studies have utilized such approaches to determine PS expression on bacterial cells (24, 25). In fact, leaflets of outer membranes from Gram-negative bacteria are highly asymmetric and considered depleted of many phospho-lipids but enriched in LPS, suggesting limited accessibility to detect phospho-lipids on rickettsial membranes (26). To overcome such limitation, we pre-treated partially purified rickettsiae with ethanol and lysosome prior to PS-staining, as described elsewhere (24). We evaluated the expression of PS of *R. typhi*, *R. rickettsii*, and *R. montanensis* by flow cytometry using APC-conjugated AnxV and showed that all three rickettsiae express PS on the surface (Figs. 3A-C). To address the PS expression within each *Rickettsia* species, we employed IFA as described previously (20, 22), and showed that anti-PS antibody-stained *R. typhi*, *R. rickettsii*, or *R. montanensis* express significant levels of PS when compared to IgG-antibody-stained bacteria (Figs. 3D-G, and Fig. S2). These data suggest that pathogenic and non-pathogenic *Rickettsia* species express external and internal PS moieties. As our bacterial inocula were purified using the glass bead method (27), we were wondering whether the observed PS-mediated phagocytic phenotype could be a technical artifact of the actual purification process. To investigate the extent of potential PS-carryover from host cell membrane remnants, we infected WT or CD300f^-/-^ BMDMΦ with purified rickettsiae of different purity levels [partially purified (PP), crudely purified (CP), and sucrose purified (SC)] as described in materials and methods section, and performed phagocytosis studies for various length of time. Similar to our data showing in Fig. 1 phagocytosis of *R. typhi* (PP), *R. rickettsii* (PP), and *R. montanensis* (PP) was significantly reduced in CD300f^-/-^ BMDMΦ compared to that in WT BMDMΦ (Figs. S3A, S3D, and S3G). Intriguingly, phagocytosis studies using either CP (Figs. S3B, S3E, and S3H) or SP (Figs. S3C, S3F, and S3I) rickettsiae, revealed similar internalization kinetics, suggesting that any PS originated from host cell membrane remnants did not alter the observed phagocytic process.

**Fig. 3.**
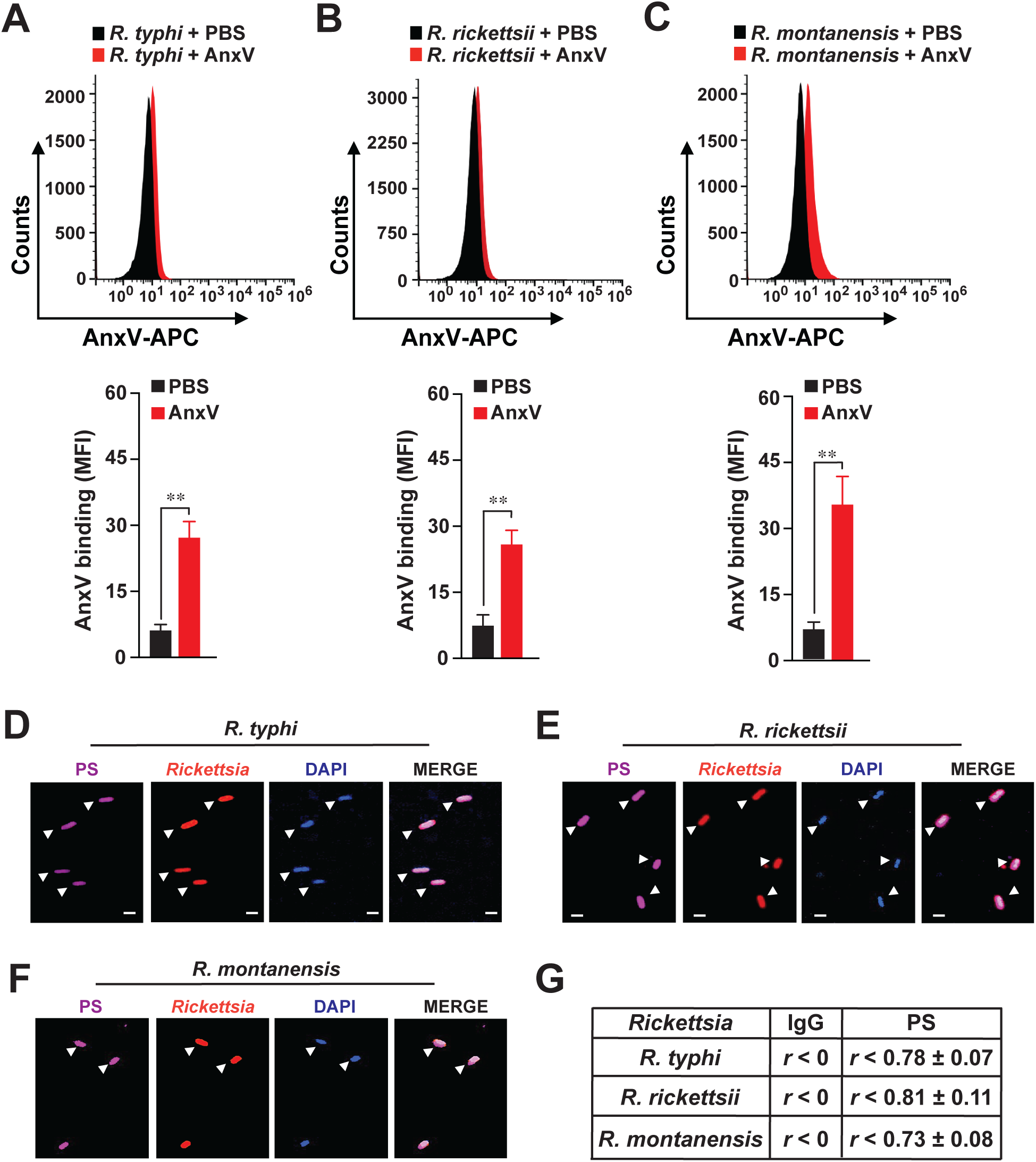
Pathogenic and non-pathogenic rickettsiae express phosphatidylserine. (A-C) Partially purified *R. typhi*, *R. rickettsii*, and *R. montanensis* were fixed with 70% ethanol and treated with lysozyme. Bacteria were then incubated with APC-conjugated AnxV for 30 min and acquired using a Becton-Dickinson Accuri C6 machine. Samples were analyzed by FlowJo software and fluorescence was expressed as mean fluorescence intensity (MFI). Error bars in panels A-C, represent means ± SEM from five independent experiments. NS, nonsignificant; \*\**P* ≤ 0.01. (D-G) Partially purified *R. typhi*, *R. rickettsii*, and *R. montanensis* were fixed with 4 % PFA and analyzed by IFA. Samples were stained with Alexa Fluor-594-conjugated anti-*Rickettsia* [SFG (1:100); TG (1:500)] Abs as well as Alexa Fluor-647-conjugated anti-PS Ab (1:100). The cell nuclei were stained with 4’,6-diamidino-2-phenylindole (DAPI). (G) Co-localization between *Rickettsia* and anti-PS or anti-IgG control Ab staining’s was analyzed using Coloc 2 plugin Fiji software and displayed as Pearson correlation coefficient (*r*) (75); 0 < *r* < 0.39 low correlation; 0.4 < *r* < 0.59 moderate correlation; 0.6 < *r* < 0.79 high correlation; 0.8 < *r* < 1 very high correlation. Bars in panels D-F, 1 μm. Approximately 100 bacteria-infected cells were analyzed per condition and time point. Presented images are representative of three independent experiments.

### CD300f exacerbates the pathogenesis of rickettsiae

Given our findings of PS-CD300f-mediated engulfment of rickettsiae in MΦ, we next tested *in vivo* the role of CD300f in *Rickettsia* colonization by employing our established C57BL/6J WT mouse model of severe (∼LD_50_) rickettsiosis (20). In agreement with our published findings, we observed a mortality rate of ∼50% in WT mice using *R. typhi* (∼68 hpi) and *R. rickettsii* (∼50 hpi), while *R. montanensis*-infected mice showed no signs of lethality (Figs. 4A, 4C, and 4E). Our findings further revealed that, *R. typhi*-, *R. rickettsii*-or *R. montanensis*-infected CD300f^-/-^ mice showed no signs of lethality during course of infection (Figs. 4A, 4C, and 4E). Bacterial burden in spleens from all three *Rickettsia*-infected WT and CD300f^-/-^ mice confirmed successful infection at day 3 post-infection (Figs. 4B, 4D, and 4F). Furthermore, bacterial burdens in the spleens of *R. typhi*-, *R. rickettsii*-, and *R. montanensis*-infected WT mice were significantly higher than that of rickettsiae-infected CD300f^-/-^ mice at day 3 and 7 (Figs. 4B, 4D, and 4F). This correlated with the differences observed in the spleen weights of the rickettsiae-infected animals (Fig. S4). These data suggest that presence of CD300f exacerbates rickettsiosis *in vivo*.

**Fig. 4.**
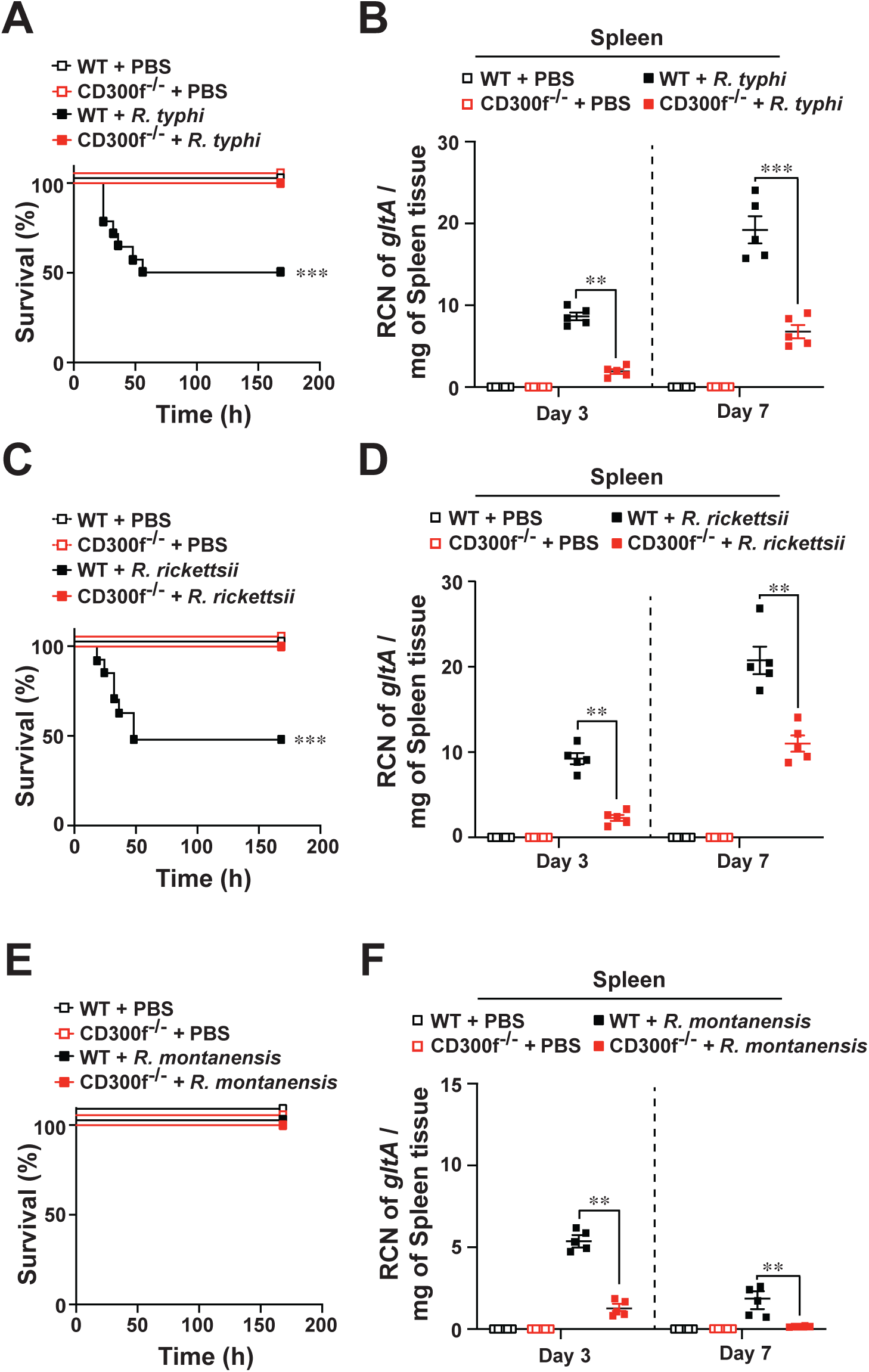
Efferocytic receptor, CD300f, exacerbates the pathogenesis of rickettsiosis. C57BL/6J WT or CD300f^-/-^ mice were injected via tail vein (*i.v.*) with *R. typhi* (A-B), *R. rickettsii* (C-D), *R. montanensis* (E-F), and PBS (10^6^ PFU, n = 12 for each treatment). Survival was monitored for 7 days (A, C, and E). Bacterial burden was tested in spleens of *R. typhi*-, *R. rickettsii*-, *R. montanensis*-, or PBS-injected WT and CD300f^-/-^ mice shown in panels B, D, F by RT-qPCR at day 3 and 7 (n = 5 for each treatment). Relative copy number (RCN) of *gltA* expression of rickettsiae was normalized by the expression of the housekeeping host gene, *GAPDH*. Error bars in panels B, D, and E represent means ± SEM from five independent experiments. \*\**P* ≤ 0.01; \*\*\**P* ≤ 0.005.

### CD300f-expressing macrophages are crucial in regulating rickettsiae invasion *in vivo*

Given the presented data and preceding findings from others and our laboratory highlighting the importance of MΦ in regulating rickettsiae infection (20, 22, 28–31), we sought to determine the contributing role of MΦ in the observed susceptibility difference between *R. typhi*-or *R.* rickettsii-infected CD300f^-/-^ and WT mice. In this effort, we focused on both pathogenic rickettsiae as infection studies using the non-pathogenic *R. montanensis* strain did not reveal any survival difference in our infection assays (Figs. 4A, 4C, and 4E). We injected C57/B6J WT and CD300f^-/-^-deficient mice via tail vein (*i.v.*) with liposome-encapsulated phosphate-buffered saline (PBS)-or dichloromethylene bisphosphonate (Cl_2_MBP) at 48 and 24 hrs to deplete endogenous MΦ as described previously (18, 20). Next, mice were infected with a low dose (10^5^ PFU) of either *R. typhi* or *R. rickettsii*, which resulted in the development of a mild form of rickettsiosis (∼LD_25_) in WT mice (20) (Figs. 5A, 5B, and 5E). Our data further revealed that depletion of MΦ not only enhanced the mortality rate of *R. typhi*- and *R. rickettsii-*infected WT mice but also impaired the survival advantage observed in CD300f^-/-^ mice (Figs. 5B, and 5E). In contrast, injection of PBS-liposomes did not alter the mortality rate of *R. typhi*- or *R. rickettsii*-infected WT or CD300f^-/-^ mice (Figs. 5B, and 5E). Of note, development of splenomegaly (represented by an increase in spleen weight) and consequently bacterial loads within the spleens were significantly elevated in rickettsiae-infected Cl_2_MBP-treated mice as compared to that of bacteria-infected PBS-liposomes treated WT and CD300f^-/-^ animals (Figs. 5C-D, and 5F-G). Taken together our data suggest that the presence of CD300f-expressing MΦ plays a critical role in regulating rickettsial infection *in vivo*.

**Fig. 5.**
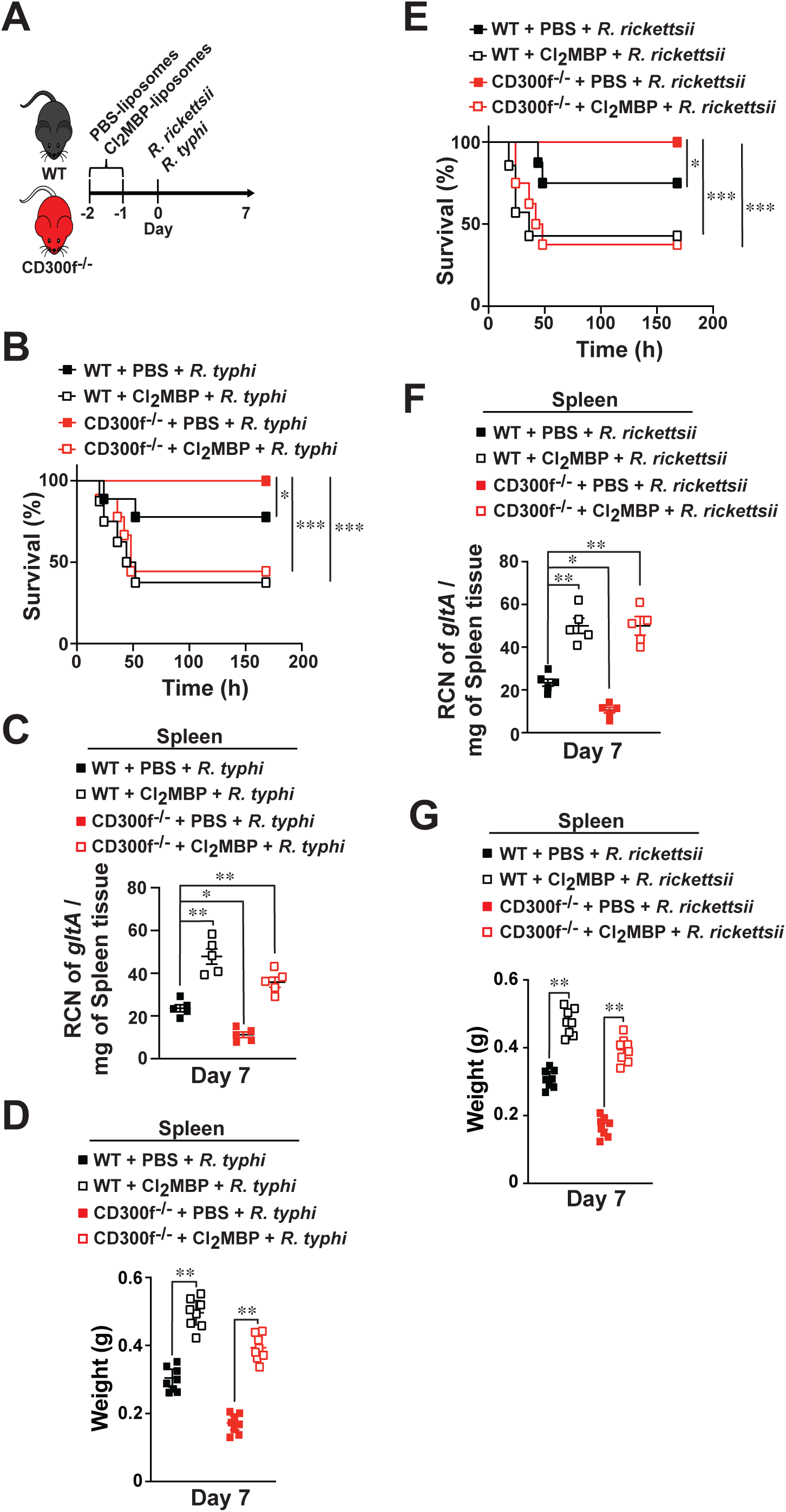
CD300f-expressing macrophages are critical in regulating rickettsiae infection. (A) C57/B6J WT or CD300f^-/-^-deficient mice (n = 12 per experimental group) were *i.v.* injected twice (48 and 24 hrs) with PBS- or Cl_2_MBP-liposomes (200 μl/per mouse) followed by *i.v.* injections with 10^5^ PFU of *R. typhi* (B-D), or *R. rickettsii* (E-G). Survival was monitored for 7 days (B and E). Bacterial burden (C and F) was determined in spleens of *Rickettsia*-injected WT or CD300f^-/-^ mice by RT-qPCR at day 7 (n = 5 for each treatment). RCN of *gltA* expression of rickettsiae was normalized by the expression of the housekeeping host gene, *GAPDH*. Spleen weights (D and G) from injected animals were evaluated at day 7 day (n = 8). Error bars in panels C, D, F, and G represent means ± SEM from five independent experiments; \**P* ≤ 0.05; \*\**P* ≤ 0.01; \*\*\**P* ≤ 0.005.

### CD300f on macrophages regulates rickettsiae pathogenesis *in vivo*

To further assess the role of CD300f-expressing MΦ in modulating rickettsiosis, we injected WT mice via tail vein (*i.v.*) with PBS- or Cl_2_MBP-liposome at 72 and 48 hrs prior to injections with either WT or CD300f^-/-^ BMDMΦ at a concentration of 5 x 10^6^ cells per mouse (Fig. S5A). Twenty-four hours later, mice were injected (*i.v.*) with either *R. typhi* or *R. rickettsii* [10^5^ PFU] (Figs. S5B, and S5E) to induce mild rickettsiosis (20).

Our data showed that removal of MΦ not only enhanced the mortality rate of both *R. typhi*- and *R. rickettsii*-infected WT mice but also revealed that adoptively transferred BMDMΦ from WT mice partially rescued fatal rickettsiosis in Cl_2_MBP-treated WT mice (Figs. S5B, and S5E). In addition, injection of BMDMΦ from CD300f^-/-^ mice into Cl_2_MBP-treated WT mice further reduced the mortality rate of *R. typhi*- or *R. rickettsii*-infected WT mice to levels seen in bacterial-infected PBS-liposomes treated WT mice (Figs. S5B, and S5E). Accordingly, the development of splenomegaly and bacterial loads within the spleens was significantly increased in bacteria-infected Cl_2_MBP-treated WT mice as compared to PBS-liposomes treated WT mice (Fig. S5C-D, and S5F-G). Adoptive transfer of WT BMDMΦ reduced the spleen weights and bacterial burdens in rickettsiae-infected Cl_2_MBP-treated WT mice (Figs. S5C-D, and S5F-G). Moreover, injection of BMDMΦ from CD300f^-/-^ mice further reduced the spleen weights and bacterial burdens in Cl_2_MBP-treated WT mice, reaching to the levels observed in rickettsiae-infected PBS-liposomes treated WT mice (Figs. S5C-D and S5F-G). Of note, our adoptive transfer experiments did not completely recapitulate the protective phenotype observed in CD300f^-/-^ mice, which could be the result of not fully reestablishing the levels of endogenously expressed MΦ.

Collectively, these data suggest that rickettsiae utilize the PS-CD300f receptor axis to gain access to MΦ and further indicate that CD300f-expressing MΦ are crucial in regulating rickettsiae pathogenesis.

## Discussion

Successful host invasion by intracellular bacteria requires receptor-ligand engagement and subversion of host endocytic bactericidal pathways (32, 33). However, research on how *Rickettsia* species invade, and avoid cytosolic defense surveillance for host colonization is only now emerging beyond the difficulties posed by their obligate intracellular replication and a life cycle that involves arthropod and vertebrate hosts (4, 5). Intriguingly, *R. rickettsii* OmpA (34) does not impact the virulence in animal models, while *R. parkeri* OmpA (35) and Sca2 (36) were shown to impact virulence. Of note, the role of *R. conorii* LPS in influencing virulence remains to be resolved (37, 38). Taken collectively, these reports suggest the presence of alternative or other available membrane components that could function as ligands for host receptors. In addition, preceding research on endothelial cells, originally considered the main host target of rickettsiae, resulted in the identification of four probable host cell receptors [Ku70 (6), α2β1 (7), FGFR (8), Epac (9)]. However, selective targeting of those receptors, showed only moderate inhibition of *Rickettsia* invasion, indicating cooperative and/or alternative receptor-ligand systems for bacteria engulfment. Furthermore, no receptor-ligand mechanism for rickettsial invasion has been identified for professional phagocytes, like MΦ. To address this knowledge gap, we focused on receptor-ligand systems involved in efferocytosis, a mechanism required for cellular homeostasis, which is often hijacked by pathogens, including viruses and bacteria, to promote their host invasion (10–13, 33, 39). As this process involves the recognition of “eat me” signals [e.g., phosphatidylserine (PS)] on apoptotic cells through various PS-receptors, including MerTK, TIM1/4, or CD300s (e.g., CD300f) (10–13, 33, 39), we took advantage of the well-established *CD300f* knockout mouse model (17). In this study, we demonstrated that CD300f^-/-^ but not WT mice were protected against *R. typhi*- or *R. rickettsii*-induced fatal rickettsiosis. As preceding reports from other and our laboratory suggest that rickettsiae exhibit species-specific immune modulatory responses in MΦ to establish an intracellular niche (20, 22, 28, 30, 31, 40–45), we next evaluated the role of CD300f on MΦ during rickettsial infection. Using *in vivo* depletion and adoptive transfer studies we showed that CD300f-expressing MΦ are crucial in controlling rickettsial infection *in vivo*. Of note, our study did not evaluate a potential contributing role of other CD300f-expressing cell types, including among other neutrophils or dendritic cells. CD300f has been shown to suppress the antimicrobial activity of neutrophils against *Pseudomonas aeruginosa* and *Candida albicans* infection (46). However, infection studies using pathogenic rickettsiae suggest that neutrophils do not alter the course of rickettsiosis or contribute to the restriction of bacterial growth (43). Dendritic cells seem to mount different immune responses against *R. conorii* infection dependent on the *in vivo* model (protective type 1 response in C57BL/6 mice vs. suppressive adaptive immunity in C3H/HeN animals) (47). Intriguingly, reports from other laboratory imply that in contrast to CD300f function on MΦ, its expression inhibited efferocytosis by dendritic cells (48) suggesting that CD300f might modulates rickettsiae infection and other cellular responses differently in dendritic cells as compared to MΦ, however the precise mechanism remains to be investigated.

As the CD300 family is composed of several members, we evaluated the expression levels of *CD300a, CD300b*, *CD300c*, *CD300d*, *and CD300f* in BMDMΦ from both WT and CD300f^-/-^ mice and demonstrated that the lack of CD300f expression did not affect the expression of any of the tested CD300s. In addition, we performed invasion assays using WT, CD300f^-/-^-, and CD300d^-/-^-deficient BMDMΦ and showed that CD300f, and not CD300d, affected the internalization of *R. typhi*, *R. rickettsii*, and *R. montanensis*. Collectively, our findings indicate that CD300f, but not CD300d, plays a predominant rule in facilitating rickettsiae invasion in MΦ. However, it is important to note that MΦ express additional PS-binding receptors, including among others Ku70 (49), CD300b (18, 23), Tim4 (50), and BAI1 (51), so it is plausible that CD300f can directly or indirectly cooperate with other receptors to modulate phagocytosis of their cargo, a mechanism described for Tim4 (52) and our future work will address these questions.

As CD300f-receptor engagement by rickettsiae would likely involve a ligand, we searched for putative ligands on the rickettsial outer membrane and found the presence of several glycerophospholipids (GPLs) (53, 54), including among others phosphatidylserine (PS), and phosphatidylglycerol (PG), known “eat-me” signals involved in efferocytosis (10–12, 39). Preceding findings by others and our work described that obligate intracellular bacteria, like *Rickettsia* species, possess most enzymes required for biosynthesis of glycerophospholipids, like PS, however with some metabolic holes, because of missing enzymes in the biosynthetic pathway, illustrating the dependence on host precursor to synthesize downstream metabolites (55–57). Based on these findings we hypothesized that *Rickettsia* engulfment is mediated by a CD300f-PS systems generally used for efferocytosis. We assessed the importance of bacterial PS for engulfment in WT BMDMΦ by pre-incubating *R. typhi*, *R. rickettsii*, and *R. montanensis* with increasing concentrations of recombinant Annexin V (rAnxV), a molecule known to block PS-mediated efferocytosis (16, 17), prior to infection and showed that rAnxV reduced the engulfment of all three *Rickettsia* species in a dose-dependent manner. We started elucidating the mechanism of PS-CD300f-mediated engulfment of rickettsiae further by determining the expression of PS on the surface of *R. typhi*, *R. rickettsii*, and *R. montanensis* using flow cytometry in combination with fluorescently labeled AnxV. Our data showed that all three rickettsiae express PS on the surface. In addition, employing IFA using Alexa Fluor 647-conjugated anti-PS antibody staining we demonstrated the PS expression in *R. typhi*, *R. rickettsii*, and *R. montanensis*, indicating that pathogenic and non-pathogenic rickettsiae express external and internal PS moieties. We further addressed the potential PS-carryover from host cell membrane remnants, by performing invasion assays of all three *Rickettsia,* using purified bacteria with different purities [partially purified (PP), crudely purified (CP), and sucrose purified (SC)], and showed a similar internalization kinetics among all samples, suggesting that any potential PS-carryover was not a contributing factor to the phagocytic process. However, it is important to note that not all externalized PS is functionally equivalent, and therefore it is still feasible that the PS found on the bacterial outer membrane is not directly involved in the phagocytosis process directly, which was described for other cells [(e.g., monocytes and macrophages (58, 59)]. It also remains plausible that the PS promoting *Rickettsia* engulfment is provided by a bystander apoptotic cell remnant, a mechanism described for *Trypanosoma brucei* (60), or via a PS-cloaking mechanism a pathway described for *Listeria monocytogenes* (61). Although PS is considered as a key physiological ligand for both human and mouse CD300f receptors (62–65), recent findings indicate a subtle differences in ligand specificity dependent on the cellular context (66, 67). Thus, it is tempting to speculate that other ligands may contribute to the internalization process of rickettsiae, which will be addressed in our future work.

In sum, we now present a working model for both pathogenic and non-pathogenic rickettsiae that utilize bacterial-PS to bind to the efferocytic CD300f receptor on MΦ to facilitate host invasion.

## Materials and Methods

### Animals

All experiments were conducted in fully AAALAC-accredited program using 8-to 10-week-old female C57BL/6J wild-type (WT), CD300d^−/−^, or CD300f^−/−^ mice in a specific-pathogen-free environment according to the University of Maryland School of Medicine Institutional Animal Care and Use Committee (IACUC protocol: AUP-00000110).

### Antibodies and Reagents

Antibodies against whole *Rickettsia* (SFG or TG) were raised in house (20, 22, 45). Lysozyme and anti-phosphatidylserine (PS) antibody were purchased from Sigma. ProLong Gold antifade mounting medium with DAPI (4′,6-diamidino-2-phenylindole), paraformaldehyde (PFA), and Alexa 594, 647-conjugated secondary antibodies as well as Alexa Fluor 488-conjugated wheat germ agglutinin (WGA) were purchased from Thermo Fisher Scientific.

### Bacterial strains, cell culture, and infection

Vero76 cells (African green monkey kidney, ATCC, RL-1587) were maintained in minimal Dulbecco’s modified Eagle’s medium (DMEM) supplemented with 10% heat-inactivated fetal bovine serum (FBS) at 37°C with 5% CO_2_. *R. rickettsii* (*Sheila Smith*) strain was obtained from Dr. Ted Hackstadt (Rocky Mountain Laboratories, NIH, MT, USA) and *R. typhi* strain (Wilmington) was obtained from CDC. All *Rickettsia* species were propagated in Vero76 cells grown in DMEM supplemented with 5% FBS at 34°C with 5% CO_2_. Rickettsiae from infected host cells were purified as described previously (21, 27, 68, 69). Briefly, rickettsiae-infected host cells were disrupted by vortexing with 1 mm glass beads. The disrupted host cells were centrifuged at 250 x *g* for 5 min at 4°C to remove host cell debris or any remaining intact host cells and collected the supernatant carrying released crudely purified (CP) rickettsiae. The supernatant, carrying released rickettsiae (CP), was then centrifuged at 9,000 x *g* for 3 min at 4°C. The pellet was collected, and washed in 1 x PBS, to obtain partially purified (PP) rickettsiae. The rickettsiae (PP) were resuspended in 1 x PBS and layered on a 20% sucrose cushion at a 1:1 ratio and centrifuged at 16,000 x *g* for 15 min at 4°C. The pellet was collected and washed in 1 x PBS to obtain sucrose purified (SP) rickettsiae. For early stages of infection [before the doubling time (8 to 10 hrs) of rickettsiae], a higher multiplicity of infection (MOI) of 20 [*e.g.*, 0.5, and 2 hrs post-infection (hpi) for Immunofluorescent assay] was used to ensure the presence of sufficient number of bacteria, as compared to MOI of 5 at later time points (24 hpi), to determine the biological functions of the bacteria during host infection (21, 22, 45, 70, 71). Tail vein injection (*i.v.*) of partially purified (PP) *Rickettsia* (10^5^-10^6^ PFU), resuspended in PBS was used to initiate infection in mice as described previously (20). Splenic tissue specimens were collected at the indicated times and used for bacterial burden analysis by RT-qPCR described below.

### Differentiation of bone marrow-derived macrophages

Bone marrow cells were isolated from femurs and tibias of WT, CD300d^-/-^, or CD300f^-/-^ mice. Differentiation was induced by culturing bone marrow cells in RPMI 1640 medium supplemented with 10% FBS and 30% L929-conditioned medium (a source of macrophage colony stimulating factor) and cultured for 7 days as described previously (18, 20, 22).

### RNA isolation and quantitative PCR

To determine viable bacterial number during the course of host infection, we performed RT-qPCR assay on isolated RNA (70, 72, 73). BMDMΦ samples were collected at different times of post infection, while spleens were collected at day 7 post infection. RNA was extracted from 1 x 10^6^ BMDMs or 100 μl of organ homogenate using the Quick-RNA miniprep kit (ZymoResearch). The iScript Reverse Transcription Supermix kit (Bio-Rad) was used to synthesize cDNAs from 200 ng of RNA according to the manufacturer’s instructions. The qPCR amplification and detection were performed on an QuantStudio™ 3 Real-Time PCR Systems (Applied Biosystem by Thermo Fischer) as described previously (20, 22, 74). Briefly, qPCR was carried out using SYBR Green (Thermo Fisher Scientific), 2 μl cDNA and 1 μM each of the following oligonucleotides for rickettsial (housekeeping) citrate synthase gene (*gltA*), and host (housekeeping) *GAPDH* gene. Oligonucleotides for detecting murine *CD300a*, *CD300b*, *CD300c*, *CD300d*, and *CD300f* were obtained from Qiagen. Cycling conditions were as follows: 1 cycle at 95°C for 3 min; 40 cycles at 95°C for 15 sec, 55°C for 15 sec, and 72°C for 20 sec; and 1 cycle to generate the dissociation curve. Melting curve analyses were performed at the end of each run to ensure that only one product was amplified. Relative copy number (RCN) of *gltA* expression was normalized by the expression of the *GAPDH* and calculated with the equation: RCN = E^−ΔCt^, where E = efficiency of PCR, and Ct = Ct *target* − Ct *GAPDH* as described previously (20, 22).

### *In vivo* depletion of macrophages following rickettsiae challenge

C57/B6J WT or CD300f^-/-^ mice were injected via tail vein (*i.v.*) using liposome-encapsulated PBS-or dichloromethylene bisphosphonate (Cl_2_MBP), as described previously (18, 20). Mice were injected twice (48 and 24 hrs) with PBS-or Cl_2_MBP-liposomes (200 µl/per mouse) followed by injections (*i.v.*) twenty-four hours later with 10^5^ PFU of *R. typhi* or *R. rickettsii*. Survival was monitored for 7 days. Bacterial burden was determined in spleens of uninfected or *R. typhi*-and *R. rickettsii*-injected mice by RT-qPCR at day 7 (n = 5 for each treatment).

### Adoptive transfer of bone marrow derived macrophages following *Rickettsia* challenge

C57/B6J WT mice were injected via tail vein (*i.v.*). Mice were injected twice (72 and 48 hrs) with PBS-or Cl_2_MBP-liposomes (200 µl/per mouse) followed by injections with BMDMΦ (5 x 10^6^ cells/mouse) from CD300f^-/-^, or WT mice, as described previously (18, 20). Twenty-four hours later, mice were injected (*i.v.*) with 10^5^ PFU of *R. typhi* or *R. rickettsii*. Survival was monitored for 7 days, and bacterial burden was determined in spleens of uninfected or *Rickettsia*-injected mice by RT-qPCR.

### Immunofluorescent assay (IFA)

Eight-well chamber slides were seeded with WT, CD300d^-/-^, or CD300f^-/-^ BMDMΦ (∼ 50 x 10^4^ cells/well) and infected using purified *Rickettsia* species (MOI = 20 [0.5 and 2 hrs] or 5 [24 hrs]) as described previously (20–22, 45). Briefly, rickettsiae were added to BMDMΦ and incubated for various length of time at 34°C. Following incubation, cells were washed three times with 1 x PBS and fixed with 4% paraformaldehyde (PFA) for 20 min at room temperature. For immunofluorescent assay (IFA), cells were stained with the following primary Abs: anti-*Rickettsia* [SFG (1:100 dilution); TG (1:500 dilution)], anti-PS or anti-IgG control Abs (1:100 dilution), and Alexa Fluor 488-conjugated WGA, as described previously (21). Cells were then washed with 1 x PBS and incubated for 1 h with anti-Alexa Fluor 594 and anti-Alexa Fluor 647 Abs diluted 1:1,500 in Ab-dilution buffer. Next, cells were washed with 1 x PBS and mounted with ProLong Gold antifade mounting medium containing DAPI. Images were acquired using the Nikon W-1 spinning disk confocal microscope (University of Maryland Baltimore, Confocal Core Facility). Co-localization strength between *Rickettsia*, and phosphatidylserine (PS) was analyzed using Fiji software as described previously (20, 22, 45). The percentage of internalized bacteria (approximately 200 bacteria were counted per strain and time point) was calculated by dividing the number of extracellular bacteria by the total number of bacteria, multiplying by 100, and then subtracting this number from 100% to get the percentage of intracellular bacteria.

For infection assays using recombinant Annexin V protein (rAnxV), partially purified *Rickettsia* were incubated with various concentrations of regular or heat inactivated rAnxV for 0.5 h at 4°C. Rickettsiae treated with rAnxV were used to infect BMDMΦ for additional 2 h at 34°C. Following incubation, cells were washed, fixed, and stained with anti-*Rickettsia* Abs as described above.

### Detection of phosphatidylserine by flow cytometry

Partially purified rickettsiae (∼ 1 x 10^7^) were fixed with in 70% ethanol (vol/vol) under constant mixing. Bacteria were centrifuged and pellet was resuspended in 1 x PBS containing lysozyme (100 mg/L). Samples were incubated for 1 h at 37°C and filtered using a nylon mesh. Bacteria were stained using an Annexin V apoptosis detection kit (BioLegend) following the manufactures instructions. Briefly, bacteria were resuspended in an AnxV-dilution buffer at 1:10 and incubated with APC-conjugated Annexin V for 30 min in the dark on ice. Samples were washed, resuspended in 1 x PBS buffer and acquired using a Becton-Dickinson Accuri C6 machine. Samples were analyzed by FlowJo software (version 10).

### Statistical analysis

Data sets were considered statistically significant when a *P* ≤ 0.05 value was obtained by unpaired Student’s *t*-test (two-tailed), paired Student’s *t*-test (two-tailed), a one-way ANOVA with either multiple comparisons or comparison to WT bacteria, a two-way ANOVA, or a log-rank (Mantel-Cox) test. Statistical analyses were performed using GraphPad Prism Software, version 8 and samples were denoted using the following asterisks: \**P*≤ 0.05; \*\**P* ≤ 0.01; \*\*\**P* ≤ 0.005; \*\*\*\**P* ≤ 0.001. Data are presented as mean ± SEM, unless stated otherwise.

## Acknowledgments

We are grateful to Ted Hackstadt (Rocky Mountain Laboratories, NIH, MT, USA) for generously providing us with essential biological reagents, including antibodies and *Rickettsia* strains. We further would like to thank Abdu F. Azad and Magda Sexton (University of Maryland School of Medicine, Baltimore, MD, USA) for their support and guidance during design and planning of this CD300f project.

This work was supported with funds from the NIAID/NIH grants (R01AI017828 and R01AI126853 to M.S.R., and R21AI166821 to O.H.V. and M.S.R.).

## Supplemental figure legends

**Fig. S1. CD300f, is highly expressed and plays a key role on macrophages to modulate rickettsiae invasion *in vitro*.**

(A) Expression levels of murine *CD300a*, *CD300b*, *CD300c*, *CD300d*, *CD300f* were assessed in uninfected WT and CD300f^-/-^ BMDMΦ by RT-qPCR (n = 3 per group). Expression levels of all *CD300s* was normalized by *GAPDH* expression. (B-D) Rickettsial invasion was monitored by IFA at 2 hpi as described previously (21). (E-G) Bacterial burden in WT and CD300f^-/-^ BMDMΦ infected with *R. typhi*, *R. rickettsii*, or *R. montanensis* was determined at 2 hpi (n = 5 per group) by RT-qPCR. RCN of *gltA* expression of rickettsiae was normalized by the expression of the housekeeping host gene, *GAPDH*. Error bars in panels A-G represent means ± SEM from 3-5 independent experiments. NS, nonsignificant; \**P* ≤ 0.05; \*\**P* ≤ 0.01.

**Fig. S2. Anti-phosphatidylserine antibody specificity staining in purified rickettsiae.**

Partially purified *R. typhi*, *R. rickettsii*, and *R. montanensis* were fixed with 4 % PFA and stained with Alexa Fluor-594-conjugated anti-*Rickettsia* [SFG (1:100); TG (1:500)] Abs as well as Alexa Fluor-647-conjugated anti-IgG control Ab. The cell nuclei were stained with 4’,6-diamidino-2-phenylindole (DAPI). Co-localization between *Rickettsia* and anti-IgG antibody staining was analyzed using Coloc 2 plugin Fiji software. Bars in panels A-C, 1 μm. Approximately 100 bacteria-infected cells were analyzed per condition and time point. Presented images are representative of three independent experiments.

**Fig. S3. Assessment of PS-carryover from different *Rickettsia*-Host cell preparations.**

Three different types of inocula [partially purified (PP), crudely purified (CP), and sucrose purified (SP)] from *R. typhi* (A-C), *R. rickettsii* (D-F), or *R. montanensis* (H-I) were utilized to infect WT or CD300f^-/-^ BMDMΦ for 2 and 24 hpi using a MOI of 20 (2 hpi) and 5 (24 hpi) respectively. Rickettsial invasion was monitored by IFA as described previously (21). Error bars in panels A-I represent means ± SEM from 5 independent experiments. NS, nonsignificant; \*\**P* ≤ 0.01.

**Fig. S4. Splenic data during infection of pathogenic and non-pathogenic *Rickettsia* species.** C57BL/6J WT and CD300f^-/-^ mice injected (*i.v.*) with *R. typhi*, *R. rickettsii*, *R. montanensis*, and PBS (dose of 10^6^ PFU). Spleen weights from injected animals were evaluated at day 7 day (n = 8). Error bars represent means ± SEM from five independent experiments. NS, nonsignificant; \**P* ≤ 0.05; \*\*\*\**P* ≤ 0.001.

**Fig. S5. CD300f-expressing macrophages modulate rickettsiae pathogenesis.** (A) C57/B6J WT mice (n = 12 per experimental group) were injected via tail vein (*i.v.*) using liposome-encapsulated PBS-or dichloromethylene bisphosphonate (Cl_2_MBP). Mice were injected twice (72 and 48 hrs) with PBS-or Cl_2_MBP-liposomes (200 μl/per mouse) followed by injections with BMDMΦ (5 x 10^6^ cells/mouse) from CD300f^-/-^, or WT mice. Twenty-four hours later, mice were injected (*i.v.*) with 10^5^ PFU of *R. typhi* (B-D) or *R. rickettsii* (E-G). Survival was monitored for 7 days (B and E). Bacterial burden (C and F) was determined in spleens of *Rickettsia*-injected WT or CD300f^-/-^ mice by RT-qPCR at day 7 (n = 5 for each treatment). RCN of *gltA* expression of rickettsiae was normalized by the expression of the housekeeping host gene, *GAPDH*. Spleen weights (D and G) from injected animals were evaluated at day 7 day (n = 8). Error bars in panels C, D, F, and G represent means ± SEM from five independent experiments; NS, non-significant; \**P* ≤ 0.05; \*\**P* ≤ 0.01; \*\*\**P* ≤ 0.005.

## References

1. Drexler NA, Yaglom H, Casal M, Fierro M, Kriner P, Murphy B, Kjemtrup A, Paddock CD. 2017. Fatal Rocky Mountain Spotted Fever along the United States-Mexico Border, 2013-2016. Emerg Infect Dis 23:1621–1626.

2. Billeter SA, Metzger ME. 2017. Limited Evidence for Rickettsia felis as a Cause of Zoonotic Flea-Borne Rickettsiosis in Southern California. J Med Entomol 54:4–7.

3. Blanton LS, Idowu BM, Tatsch TN, Henderson JM, Bouyer DH, Walker DH. 2016. Opossums and cat fleas: New insights in the ecology of murine typhus in Galveston, Texas. American Journal of Tropical Medicine and Hygiene 95:457–461.

4. Sahni A, Fang R, Sahni SK, Walker DH. 2019. Pathogenesis of Rickettsial Diseases: Pathogenic and Immune Mechanisms of an Endotheliotropic Infection. Annu Rev Pathol 14:127–152.

5. Voss OH, Rahman MS. 2021. Rickettsia-host interaction: strategies of intracytosolic host colonization. Pathog Dis 79.

6. Martinez JJ, Seveau S, Veiga E, Matsuyama S, Cossart P. 2005. Ku70, a component of DNA-dependent protein kinase, is a mammalian receptor for Rickettsia conorii. Cell 123:1013–1023.

7. Hillman RD, Baktash YM, Martinez JJ. 2013. OmpA-mediated rickettsial adherence to and invasion of human endothelial cells is dependent upon interaction with α2β1 integrin. Cell Microbiol 15:727–741.

8. Sahni A, Patel J, Narra HP, Schroeder CLC, Walker DH, Sahni SK. 2017. Fibroblast growth factor receptor-1 mediates internalization of pathogenic spotted fever rickettsiae into host endothelium. PLoS One 12.

9. Gong B, Shelite T, Mei FC, Ha T, Hu Y, Xu G, Chang Q, Wakamiya M, Ksiazek TG, Boor PJ, Bouyer DH, Popov VL, Chen J, Walker DH, Cheng X. 2013. Exchange protein directly activated by cAMP plays a critical role in bacterial invasion during fatal rickettsioses. Proceedings of the National Academy of Sciences 110:19615–19620.

10. Flannagan RS, Jaumouillé V, Grinstein S. 2012. The Cell Biology of Phagocytosis. Annual Review of Pathology: Mechanisms of Disease 7:61–98.

11. Voss OH, Tian L, Murakami Y, Coligan JE, Krzewski K. 2015. Emerging role of CD300 receptors in regulating myeloid cell efferocytosis. Mol Cell Oncol 10.4161/23723548.2014.964625.

12. Birge RB, Boeltz S, Kumar S, Carlson J, Wanderley J, Calianese D, Barcinski M, Brekken RA, Huang X, Hutchins JT, Freimark B, Empig C, Mercer J, Schroit AJ, Schett G, Herrmann M. 2016. Phosphatidylserine is a global immunosuppressive signal in efferocytosis, infectious disease, and cancer. Cell Death Differ 23:962–978.

13. Vitallé J, Terrén I, Orrantia A, Zenarruzabeitia O, Borrego F. 2019. CD300 receptor family in viral infections. Eur J Immunol 49:364–374.

14. Clark GJ, Ju X, Tate C, Hart DNJ. 2009. The CD300 family of molecules are evolutionarily significant regulators of leukocyte functions. Trends Immunol 30:209–217.

15. Borrego F. 2013. The CD300 molecules: An emerging family of regulators of the immune system. Blood 10.1182/blood-2012-09-435057.

16. Choi S-C, Simhadri VR, Tian L, Gil-Krzewska A, Krzewski K, Borrego F, Coligan JE. 2011. Cutting Edge: Mouse CD300f (CMRF-35–Like Molecule-1) Recognizes Outer Membrane-Exposed Phosphatidylserine and Can Promote Phagocytosis. The Journal of Immunology 10.4049/jimmunol.1101549.

17. Tian L, Choi SC, Murakami Y, Allen J, Morse HC, Qi CF, Krzewski K, Coligan JE. 2014. P85α recruitment by the CD300f phosphatidylserine receptor mediates apoptotic cell clearance required for autoimmunity suppression. Nat Commun 10.1038/ncomms4146.

18. Voss OH, Murakami Y, Pena MY, Lee H-N, Tian L, Margulies DH, Street JM, Yuen PST, Qi C-F, Krzewski K, Coligan JE. 2016. Lipopolysaccharide-Induced CD300b Receptor Binding to Toll-like Receptor 4 Alters Signaling to Drive Cytokine Responses that Enhance Septic Shock. Immunity 44:1365–1378.

19. Lee HN, Tian L, Bouladoux N, Davis J, Quinones M, Belkaid Y, Coligan JE, Krzewski K. 2017. Dendritic cells expressing immunoreceptor CD300f are critical for controlling chronic gut inflammation. Journal of Clinical Investigation 10.1172/JCI89531.

20. Voss OH, Cobb J, Gaytan H, Rivera Díaz N, Sanchez R, DeTolla L, Rahman MS, Azad AF. 2022. Pathogenic, but Not Nonpathogenic, Rickettsia spp. Evade Inflammasome-Dependent IL-1 Responses To Establish an Intracytosolic Replication Niche. mBio 13.

21. Rennoll-Bankert KE, Rahman MS, Gillespie JJ, Guillotte ML, Kaur SJ, Lehman SS, Beier-Sexton M, Azad AF. 2015. Which Way In? The RalF Arf-GEF Orchestrates Rickettsia Host Cell Invasion. PLoS Pathog 11:e1005115.

22. Voss OH, Gaytan H, Ullah S, Sadik M, Moin I, Rahman MS, Azad AF. 2023. Autophagy facilitates intracellular survival of pathogenic rickettsiae in macrophages via evasion of autophagosomal maturation and reduction of microbicidal pro-inflammatory IL-1 cytokine responses. Microbiol Spectr 11:e0279123.

23. Murakami Y, Tian L, Voss OH, Margulies DH, Krzewski K, Coligan JE. 2014. CD300b regulates the phagocytosis of apoptotic cells via phosphatidylserine recognition. Cell Death Differ 10.1038/cdd.2014.86.

24. Chen B, Zhao Y, Li Z, Pan J, Wu H, Qiu G, Feng C, Wei C. 2020. Immobilization of Phosphatidylserine by Ethanol and Lysozyme on the Cell Surface for Evaluation of Apoptosis-Like Decay in Activated-Sludge Bacteria. Appl Environ Microbiol 86.

25. Dwyer DJ, Camacho DM, Kohanski MA, Callura JM, Collins JJ. 2012. Antibiotic-induced bacterial cell death exhibits physiological and biochemical hallmarks of apoptosis. Mol Cell 46:561–72.

26. Horne JE, Brockwell DJ, Radford SE. 2020. Role of the lipid bilayer in outer membrane protein folding in Gram-negative bacteria. J Biol Chem 295:10340–10367.

27. Lamason RL, Kafai NM, Welch MD. 2018. A streamlined method for transposon mutagenesis of Rickettsia parkeri yields numerous mutations that impact infection. PLoS One 13:e0197012.

28. Bechelli J, Vergara L, Smalley C, Buzhdygan TP, Bender S, Zhang W, Liu Y, Popov VL, Wang J, Garg N, Hwang S, Walker DH, Fang R. 2019. Atg5 supports Rickettsia australis infection in macrophages in vitro and in vivo. Infect Immun 87.

29. Bechelli J, Rumfield CS, Walker DH, Widen S, Khanipov K, Fang R. 2021. Subversion of Host Innate Immunity by Rickettsia australis via a Modified Autophagic Response in Macrophages. Front Immunol 0:962.

30. Burke TP, Engström P, Chavez RA, Fonbuena JA, Vance RE, Welch MD. 2020. Inflammasome-mediated antagonism of type I interferon enhances Rickettsia pathogenesis. Nat Microbiol 5:688–696.

31. Engström P, Burke TP, Mitchell G, Ingabire N, Mark KG, Golovkine G, Iavarone AT, Rape M, Cox JS, Welch MD. 2019. Evasion of autophagy mediated by Rickettsia surface protein OmpB is critical for virulence. Nat Microbiol 4:2538–2551.

32. Personnic N, Bärlocher K, Finsel I, Hilbi H. 2016. Subversion of Retrograde Trafficking by Translocated Pathogen Effectors. Trends Microbiol 24:450–462.

33. Ray K, Marteyn B, Sansonetti PJ, Tang CM. 2009. Life on the inside: the intracellular lifestyle of cytosolic bacteria. Nat Rev Microbiol 7:333–340.

34. Noriea NF, Clark TR, Hackstadt T. 2015. Targeted knockout of the Rickettsia rickettsii OmpA surface antigen does not diminish virulence in a mammalian model system. mBio 6.

35. Burke TP, Engström P, Tran CJ, Langohr IM, Glasner DR, Espinosa DA, Harris E, Welch MD. 2021. Interferon receptor-deficient mice are susceptible to eschar-associated rickettsiosis. Elife 10.

36. Kleba B, Clark TR, Lutter EI, Ellison DW, Hackstadt T. 2010. Disruption of the Rickettsia rickettsii Sca2 autotransporter inhibits actin-based motility. Infect Immun2010/03/03. 78:2240–2247.

37. Feng H-MM, Whitworth T, Olano JP, Popov VL, Walker DH. 2004. Fc-dependent polyclonal antibodies and antibodies to outer membrane proteins A and B, but not to lipopolysaccharide, protect SCID mice against fatal Rickettsia conorii infection. Infect Immun 72:2222–2228.

38. Kim HK, Premaratna R, Missiakas DM, Schneewind O. 2019. Rickettsia conorii O antigen is the target of bactericidal Weil-Felix antibodies. Proc Natl Acad Sci U S A 116:19659–19664.

39. Ravichandran KS. 2010. Find-me and eat-me signals in apoptotic cell clearance: progress and conundrums. J Exp Med 207:1807–1817.

40. Borgo GM, Burke TP, Tran CJ, Lo NTN, Engström P, Welch MD. 2022. A patatin-like phospholipase mediates Rickettsia parkeri escape from host membranes. Nat Commun 13.

41. Rumfield C, Hyseni I, McBride JW, Walker DH, Fang R. 2020. Activation of ASC inflammasome driven by toll-like receptor 4 contributes to host immunity against rickettsial infection. Infect Immun 88.

42. Smalley C, Bechelli J, Rockx-Brouwer D, Saito T, Azar SR, Ismail N, Walker DH, Fang R. 2016. Rickettsia australis Activates Inflammasome in Human and Murine Macrophages. PLoS One 11:e0157231.

43. Papp S, Moderzynski K, Rauch J, Heine L, Kuehl S, Richardt U, Mueller H, Fleischer B, Osterloh A. 2016. Liver Necrosis and Lethal Systemic Inflammation in a Murine Model of Rickettsia typhi Infection: Role of Neutrophils, Macrophages and NK Cells. PLoS Negl Trop Dis 10:e0004935.

44. Moderzynski K, Papp S, Rauch J, Heine L, Kuehl S, Richardt U, Fleischer B, Osterloh A. 2016. CD4+T Cells Are as Protective as CD8+T Cells against Rickettsia typhi Infection by Activating Macrophage Bactericidal Activity. PLoS Negl Trop Dis 10.

45. Voss OH, Gillespie JJ, Lehman SS, Rennoll SA, Beier-Sexton M, Rahman MS, Azad AF. 2020. Risk1, a Phosphatidylinositol 3-Kinase Effector, Promotes Rickettsia typhi Intracellular Survival. mBio 11.

46. Ueno K, Urai M, Izawa K, Otani Y, Yanagihara N, Kataoka M, Takatsuka S, Abe M, Hasegawa H, Shimizu K, Kitamura T, Kitaura J, Miyazaki Y, Kinjo Y. 2018. Mouse LIMR3/CD300f is a negative regulator of the antimicrobial activity of neutrophils. Sci Rep 8:17406.

47. Fang R, Ismail N, Soong L, Popov VL, Whitworth T, Bouyer DH, Walker DH. 2007. Differential interaction of dendritic cells with Rickettsia conorii: impact on host susceptibility to murine spotted fever rickettsiosis. Infect Immun 75:3112–23.

48. Tian L, Choi S-C, Lee H-N, Murakami Y, Qi C-F, Sengottuvelu M, Voss O, Krzewski K, Coligan JE. 2016. Enhanced efferocytosis by dendritic cells underlies memory T-cell expansion and susceptibility to autoimmune disease in CD300f-deficient mice. Cell Death Differ 23:1086–96.

49. Sun H, Li Q, Yin G, Ding X, Xie J. 2020. Ku70 and Ku80 participate in LPS-induced pro-inflammatory cytokines production in human macrophages and monocytes. Aging 12:20432–20444.

50. Miyanishi M, Tada K, Koike M, Uchiyama Y, Kitamura T, Nagata S. 2007. Identification of Tim4 as a phosphatidylserine receptor. Nature 10.1038/nature06307.

51. Park D, Tosello-Trampont A-C, Elliott MR, Lu M, Haney LB, Ma Z, Klibanov AL, Mandell JW, Ravichandran KS. 2007. BAI1 is an engulfment receptor for apoptotic cells upstream of the ELMO/Dock180/Rac module. Nature 450:430–434.

52. Nishi C, Toda S, Segawa K, Nagata S. 2014. Tim4-and MerTK-Mediated Engulfment of Apoptotic Cells by Mouse Resident Peritoneal Macrophages. Mol Cell Biol 34:1512–1520.

53. Winkler HH, Miller ET. 1978. Phospholipid composition of Rickettsia prowazeki grown in chicken embryo yolk sacs. J Bacteriol.

54. Tzianabos T, Moss CW, McDade JE. 1981. Fatty acid composition of rickettsiae. J Clin Microbiol.

55. Lin M, Grandinetti G, Hartnell LM, Bliss D, Subramaniam S, Rikihisa Y. 2020. Host membrane lipids are trafficked to membranes of intravacuolar bacterium Ehrlichia chaffeensis. Proceedings of the National Academy of Sciences 117:8032–8043.

56. V A, CA B, TP B, DK N, MD W. 2019. A Metabolic Dependency for Host Isoprenoids in the Obligate Intracellular Pathogen Rickettsia parkeri Underlies a Sensitivity to the Statin Class of Host-Targeted Therapeutics. mSphere 4.

57. Driscoll TP, Verhoeve VI, Guillotte ML, Lehman SS, Rennoll SA, Beier-Sexton M, Rahman MS, Azad AF, Gillespie JJ. 2017. Wholly *Rickettsia* ! Reconstructed Metabolic Profile of the Quintessential Bacterial Parasite of Eukaryotic Cells. mBio 8:e00859–17.

58. Callahan MK, Williamson P, Schlegel RA. 2000. Surface expression of phosphatidylserine on macrophages is required for phagocytosis of apoptotic thymocytes. Cell Death Differ 7:645–653.

59. Appelt U, Sheriff A, Gaipl US, Kalden JR, Voll RE, Herrmann M. 2005. Viable, apoptotic and necrotic monocytes expose phosphatidylserine: Cooperative binding of the ligand Annexin V to dying but not viable cells and implications for PS-dependent clearance [2]. Cell Death Differ 10.1038/sj.cdd.4401527.

60. DaMatta RA, Seabra SH, Deolindo P, Arnholdt AC V., ManhÃ£es L, Goldenberg S, de Souza W. 2007. Trypanosoma cruzi exposes phosphatidylserine as an evasion mechanism. FEMS Microbiol Lett 266:29–33.

61. Czuczman MA, Fattouh R, van Rijn JM, Canadien V, Osborne S, Muise AM, Kuchroo VK, Higgins DE, Brumell JH. 2014. Listeria monocytogenes exploits efferocytosis to promote cell-to-cell spread. Nature 509:230–234.

62. Tian L, Choi S-C, Murakami Y, Allen J, Morse HC, Qi C-F, Krzewski K, Coligan JE. 2014. p85α recruitment by the CD300f phosphatidylserine receptor mediates apoptotic cell clearance required for autoimmunity suppression. Nat Commun 5:3146.

63. Voss OH, Tian L, Murakami Y, Coligan JE, Krzewski K. 2015. Emerging role of CD300 receptors in regulating myeloid cell efferocytosis. Mol Cell Oncol 2:e964625.

64. Voss OH, Murakami Y, Pena MY, Lee H-N, Tian L, Margulies DH, Street JM, Yuen PST, Qi C-F, Krzewski K, Coligan JE. 2016. Lipopolysaccharide-Induced CD300b Receptor Binding to Toll-like Receptor 4 Alters Signaling to Drive Cytokine Responses that Enhance Septic Shock. Immunity 44:1365–78.

65. Choi S-C, Simhadri VR, Tian L, Gil-Krzewska A, Krzewski K, Borrego F, Coligan JE. 2011. Cutting edge: mouse CD300f (CMRF-35-like molecule-1) recognizes outer membrane-exposed phosphatidylserine and can promote phagocytosis. J Immunol 187:3483–7.

66. Izawa K, Isobe M, Matsukawa T, Ito S, Maehara A, Takahashi M, Yamanishi Y, Kaitani A, Oki T, Okumura K, Kitamura T, Kitaura J. 2014. Sphingomyelin and ceramide are physiological ligands for human LMIR3/CD300f, inhibiting FcεRI-mediated mast cell activation. J Allergy Clin Immunol 133:270–3.e1-7.

67. Izawa K, Maehara A, Isobe M, Yasuda Y, Urai M, Hoshino Y, Ueno K, Matsukawa T, Takahashi M, Kaitani A, Shiba E, Takamori A, Uchida S, Uchida K, Maeda K, Nakano N, Yamanishi Y, Oki T, Voehringer D, Roers A, Nakae S, Ishikawa J, Kinjo Y, Shimizu T, Ogawa H, Okumura K, Kitamura T, Kitaura J. 2017. Disrupting ceramide-CD300f interaction prevents septic peritonitis by stimulating neutrophil recruitment. Sci Rep 7:4298.

68. Kaur SJ, Rahman MS, Ammerman NC, Beier-Sexton M, Ceraul SM, Gillespie JJ, Azad AF. 2012. TolC-dependent secretion of an ankyrin repeat-containing protein of *Rickettsia typhi*. J Bacteriol 194:4920–32.

69. Lehman SS, Noriea NF, Aistleitner K, Clark TR, Dooley CA, Nair V, Kaur SJ, Rahman MS, Gillespie JJ, Azad AF, Hackstadt T. 2018. The rickettsial ankyrin repeat protein 2 is a type IV secreted effector that associates with the endoplasmic reticulum. mBio 9.

70. Rahman MS, Ammerman NC, Sears KT, Ceraul SM, Azad AF. 2010. Functional characterization of a phospholipase A(2) homolog from Rickettsia typhi. J Bacteriol 192:3294–3303.

71. Pang H, Winkler HH. 1996. Transcriptional analysis of the 16s rRNA gene in Rickettsia prowazekii. J Bacteriol 178:1750–1755.

72. Klein PG, Juneja VK. 1997. Sensitive detection of viable Listeria monocytogenes by reverse transcription-PCR. Appl Environ Microbiol 63:4441–4448.

73. Yaron S, Matthews KR. 2002. A reverse transcriptase-polymerase chain reaction assay for detection of viable Escherichia coli O157:H7: investigation of specific target genes. J Appl Microbiol 92:633–640.

74. Rennoll SA, Rennoll-Bankert KE, Guillotte ML, Lehman SS, Driscoll TP, Beier-Sexton M, Sayeedur Rahman M, Gillespie JJ, Azad AF. 2018. The cat flea (Ctenocephalides felis) immune deficiency signaling pathway regulates Rickettsia typhi infection. Infect Immun 86.

75. Dunn KW, Kamocka MM, McDonald JH. 2011. A practical guide to evaluating colocalization in biological microscopy. Am J Physiol Cell Physiol 300:C723–42.

